# Kin competition accelerates experimental range expansion in an arthropod herbivore

**DOI:** 10.1101/150011

**Authors:** Katrien Van Petegem, Felix Moerman, Maxime Dahirel, Emanuel A. Fronhofer, Martijn L. Vandegehuchte, Thomas Van Leeuwen, Nicky Wybouw, Robby Stoks, Dries Bonte

**Affiliations:** K. Van Petegem, M. Vandegehuchte, D. Bonte – Ghent University, Dept. Biology, K.L. Ledeganckstraat 35, 9000 Ghent, Belgium; F. Moerman, E.A. Fronhofer – Eawag: Swiss Federal Institute of Aquatic Science and Technology, Department of Aquatic Ecology, Überlanderstrasse 133, CH-8600 Dübendorf, Switzerland; University of Zürich, Department of Evolutionary Biology and Environmental Studies, Winterthurerstrasse 190, CH-8057 Zürich, Switzerland; M. Dahirel – Université de Rennes 1, UMR CNRS EcoBio, 263 avenue du Général Leclerc, 35042 Rennes, France; N. Wybouw, T. Van Leeuwen – Ghent University, Department of Crop Protection, Faculty of Bioscience Engineering, B-9000, Ghent, Belgium; Evolutionary Biology, IBED, University of Amsterdam, Science Park 904 – 1098 XH Amsterdam, The Netherlands; R. Stoks – University of Leuven, Dept. Biology, Deberiotstraat 32, 3000 Leuven, Belgium

**Keywords:** invasions, spatial sorting, experimental evolution, relatedness, *Tetranychus urticae*

## Abstract

With ongoing global change, life is continuously forced to move to novel areas, which leads to dynamically changing species ranges. As dispersal is central to range dynamics, factors promoting fast and distant dispersal are key to understanding and predicting species ranges. During range expansions, genetic variation is depleted at the expanding front. Such conditions should reduce evolutionary potential, while increasing kin competition. Organisms able to recognise relatives may be able to assess increased levels of relatedness at expanding range margins and to increase their dispersal in a plastic manner. Using individual-based simulations and experimental range expansions of a spider mite, we demonstrate that plastic responses to kin structure can be at least as important as evolution in driving range expansion speed. Because recognition of kin or kind is increasingly documented across the tree of life, we anticipate it to be a highly important but neglected driver of range expansions.

## Introduction

Range expansions and biological invasions have traditionally been studied from an ecological and conservation biological perspective, primarily in relation to climate change and invasive species (Keane & Smith 2002). The speed and extent of a range expansion can only be affected through a change in the following underlying life history traits: dispersal and reproduction (Fisher 1937). Unusual long-distance dispersal (fat tails of the dispersal kernel) were for instance brought forward as an explanation of fast range expansions of trees after the glacial periods (Reid’s paradox; Clark *et al.* 1998). Recently, researchers have realized that the dynamics of range expansions can be impacted by evolutionary changes via eco-evolutionary feedbacks: as the very drivers of range expansions (dispersal and reproduction) often have a genetic basis (e.g. Roff 2007; Saastamoinen *et al.* 2017) they can evolve during and because of the range expansion, thereby accelerating the expansion process (Shine *et al.* 2011). More specifically, the process of genetic assortment at expanding range borders results in the evolution of increased dispersal because highly dispersive genotypes colonize vacant habitat first (Phillips *et al.* 2010), after which assortative mating strengthens selection of traits affecting expansion (*i.e.* the Olympic village effect; Phillips 2015). The former is referred to as spatial sorting, the latter as spatial selection. In addition, systematically low densities at the leading edge may select for increased reproductive performance through *r-*selection (Burton *et al.* 2010). Insights into the eco-evolutionary dynamics of range expansions are largely derived from theory (e.g., Burton *et al.* 2010; Kubisch *et al.* 2013, 2014; Chuang & Peterson 2016), but empirical evidence of these eco-evolutionary dynamics is accumulating from experimental and observational studies on a wide array of taxa, including microbial systems (e.g., Datta *et al.* 2013; Fronhofer & Altermatt 2015), arthropods (Therry *et al.* 2014; Van Petegem *et al.* 2016a; Ochocki *et al.* 2017; Weiss-Lehman *et al.* 2017), vertebrates (Duckworth & Badyaev 2007; Shine 2012; Alex Perkins *et al.* 2013) and plants (Huang *et al.* 2015; Williams *et al.* 2016a).

Paradoxically, spatial sorting of genotypes during invasion is tightly associated with a successive loss of genetic variation due to subsequent founder effects. These founder effects render genetic drift important, and have the potential to further affect evolutionary change (Hallatschek *et al.* 2007). Loss of genetic variation may not only constrain evolutionary change but also increase local levels of genetic relatedness (Newman & Pilson 1997; Kubisch *et al.* 2013; Nadell *et al.* 2016). In many arthropods, for instance, single female colonisers found highly related populations (Dingle 1978).

Increased relatedness has a strong impact on dispersal, both in terms of evolutionary and plastic mechanisms (e.g. Bowler & Benton 2005, Ronce 2007). In general, dispersal is a spatial process leading to fitness maximisation (Bonte *et al.* 2014). It is therefore strongly context-and condition-dependent (Clobert *et al.* 2009) with high local densities typically boosting emigration rates, thereby enabling individuals to increase their short term performance by avoiding resource competition (Bowler & Benton 2005). Short-term performance is, however, only an incomplete measure of fitness as the latter also depends on aspects of relatedness, eventually determining the spread of genes within the population (Hamilton 1964). With increasing relatedness, competition among kin will become one of the major interactions, even in highly cooperative or social species (West *et al.* 2007). If populations are tightly kin-structured, emigration of individuals reduces local resource competition among kin while also providing a chance of colonising new habitat, even if individual dispersal costs are high. Kin competition (i.e., competition between genetic relatives) is therefore expected to be a strong driver of dispersal evolution by maximising inclusive fitness (Hamilton & May 1977).

Evidently, plastic adjustments of dispersal, conditional to the local level of relatedness, may be even more adaptive (Bitume *et al.* 2013). A major prerequisite for relatedness-dependent dispersal to be effective is the presence of kin recognition mechanisms that lead to kin discrimination (e.g., Waldman 1987; Blaustein *et al.* 1988; Waldman *et al.* 1988; Tang-Martinez 2001), that is, some sort of association and phenotype matching. This process involves the discrimination of traits of kin or self from traits of any other individual, either by learning or by means of recognition alleles. While kin recognition strategies based on spatial association and learning are widely documented, evidence is accumulating that kin recognition in the absence of social interactions and learning is neither uncommon in animals (e.g. Charpentier *et al.* 2010; Le Vin *et al.* 2010; Bonadonna & Sanz-Aguilar 2012; Krause *et al.* 2012; Leclaire *et al.* 2012) nor in plants (Dudley *et al.* 2013). Assuming that relatives (*kin* by descent) at range expansion fronts will be identical-by-state (*kind* by sharing identical traits), plastic increases of dispersal are anticipated to be a key driver of range expansions and may explain the paradox of fast expansions despite severe genetic diversity loss (Estoup *et al.* 2016). Relatedness-dependent dispersal is mechanistically driven by selection of kind rather than of kin (Queller 2011). We therefore use here the term relatedness-dependent dispersal as a special form of relatedness-dependent dispersal to refer to the recognition based on identity-by-state (IBS) rather than by Hamilton’s identity-by-descent (IBD) mechanisms. In organisms in which variation in dispersal and/or reproduction is primarily environmentally driven, relatedness-dependent dispersal following assessment of IBS may even be the primary driver of fast range expansions.

Several studies have used experimental range expansions to document evolutionary divergence in life history traits between core and edge populations, e.g. in protists (Fronhofer & Altermatt 2015), beetles (Ochocki *et al.* 2017; Weiss-Lehman *et al.* 2017), and plants (Williams *et al.* 2016b). These studies included reshuffling or replacing experiments to quantify the eco-evolutionary loop, that is how evolution feeds back on the range expansion dynamics. In this experimental procedure, individuals are systematically replaced by individuals from a source population or from a random patch in the experimental range expansion to avoid the evolution of traits related to spatial sorting or local adaptation while maintaining population densities, age, and sex structure constant. Such approaches have, however, the major drawback that patterns in relatedness and phenotype (state) are destroyed as well. Given the presumed relevance of kin competition for dispersal, and the central role of dispersal for range expansions, we expect that observed differences in expansion speed previously attributed to spatial sorting and selection could equally likely result from changes in relatedness and subsequent changes in relatedness-dependent dispersal.

We here set out to test the relative importance of plastic relatedness (IBS)-dependent dispersal compared to spatial sorting and selection for the dynamics of range expansions. We use the two-spotted spider mite *Tetranychus urticae* Koch (Acari, Tetranychidae) as a model organism because the impact of relatedness on dispersal kernels has been extensively studied in this species, which has been developed as a model for experimental evolution (Macke *et al.* 2011; Fronhofer *et al.* 2014; De Roissart *et al.* 2016; Van Petegem *et al.* 2016a). Spider mite life history traits, including dispersal, have a genetic basis but are also highly plastic in response to inter-and intra-generational environmental and social conditions (Magalhães *et al.* 2009; Bitume *et al.* 2011, 2014; Fronhofer *et al.* 2014; Van Petegem *et al.* 2015). Experimental work in this species has demonstrated the existence of kin discrimination which is presumably based on chemical, silk-related odors (Tien *et al.* 2011; Clotuche *et al.* 2012; Yoshioka & Yano 2014). Importantly, kin recognition has been shown to play an important role in condition-dependent dispersal (Bitume *et al.* 2013). We firstly developed a highly parameterized yet simple simulation model based on spider mite life histories and relatedness-by-IBS-dependent dispersal reaction norms to formalise our hypotheses and predictions. In order to provide empirical proof of principle, we subsequently conducted two experiments in which genetic diversity (i.e., evolutionary potential) and relatedness were manipulated to infer whether range expansion dynamics are jointly affected by spatial sorting and kin competition. This was accomplished by contrasting evolved trait divergence and the rate of range expansion in two sets of experimental range expansions that differed in the level of genetic variation and spatial structure. While a first experiment followed the earlier used replacement manipulations that eliminate both kin structure and evolution, a second experimental range expansion prevented evolution while maintaining kin structure.

## Material and Methods

### General model algorithm

The model is individual-based and simulates demographic and evolutionary processes along a one-dimensional array of patches (metapopulation). Patches contain resources, which are consumed by individuals at different rates depending on their life stage (juvenile or adult). Resources are refreshed on a weekly basis. We parameterised relatedness-dependent dispersal according to earlier research (partly published in (Bitume *et al.* 2013). Relatedness-dependent dispersal was here studied under average densities, that is, no further density-dependence was implemented. We thus assume the relatedness-dependent dispersal kernels to be relevant for the average population densities during range expansion. A detailed model description and additional results on in silico trait evolution are available in APPENDIX 1 in the Supporting Information.

Males and females of *Tetranychus urticae* differ in a number of aspects. Firstly, males are smaller when reaching the adult life stage, and hence contribute less to resource consumption. Secondly, dispersal behaviour differs between the two sexes, with adult females being the dominant dispersers, whereas juveniles and males disperse very little. Lastly, the species is characterized by a haplodiploid life cycle, where non-mated females only produce haploid male offspring, and mated females can produce both haploid male and diploid female eggs. The sex ratio of spider mites is female-biased, with approximately 0.66 males to females. For these reasons and for the sake of simplicity, we designed the model to only include female mites, where the genotype of the individual is passed on from mother to daughter. Individuals carry the following genetic traits: age at maturity, fecundity, longevity, and a locus determining a cue for kin recognition (one unique allele per individual). Mean relatedness (IBS) of an individual A in a patch X can be calculated as the number of individuals in patch X carrying the same allele for the loci linked to kin recognition as individual A, divided by the total number of individuals. The level of relatedness hence ranges from 0 (no related individuals present) to 1 (all individuals are related to individual A). After 80 time steps, concurring with 80 days in our experimental range expansions, both the spatial extent of the range expansion and the mean life history trait values at the core and edge were recorded. To this end, individuals present in the first patch of the metapopulation (core) or in the last three occupied patches (edge) were tracked (cf. the experimental part of the study) and the mean value of every life history trait was calculated and recorded.

The following scenarios were tested:

A. A treatment where dynamics include putative kin competition and evolution. In this scenario, females pass their allele values to the offspring. Mutations occur at a rate of 0.001 per locus and change the trait value to a randomly assigned value as during the initialisation phase. The genotype ID remains unchanged (relatedness (IBS) is unaffected by trait value mutation).
B. A treatment where dynamics do not include evolution, which is achieved by changing life history trait values (but not the one engaged in kin recognition) during reproduction at a mutational rate of 1. Therefore, in this scenario, all trait values are reset according to the initialisation procedure. Only the genotype is maintained, and therefore kin(d) structure and possible kin competition are not affected.
C. A treatment simulating the reshuffling of females. In this scenario, just as in the experimental procedure, adult females are replaced each week by random females from a stock population. Thus, both trait values and IBS are reset, eliminating both kin competition and evolutionary change.

## Experimental range expansions

### T. urticae strains

Several different strains of *T. urticae* were used within the current study: LS-VL, MR-VP, SR-VP, JPS, ALBINO, LONDON, and MIX. The LS-VL strain was originally collected in 2000 from rose plants in Ghent (Belgium) and since then maintained on common bean (*Phaseolus vulgaris*, variety Prélude) in a laboratory environment (Van Leeuwen *et al.* 2004). This strain is known to harbour high genetic diversity for studies of experimental evolution (Van Leeuwen *et al.* 2008; De Roissart *et al.* 2016). The MR-VP, SR-VP, JPS, ALBINO and LONDON strains, in contrast, were collected from different greenhouses and inbred by successive rounds of mother-son mating (see Díaz-Riquelme et al. 2016 for the followed procedure). The strain ALBINO was re-sequenced in the context of a genetic mapping study (Bryon et al 2017), confirming genome-wide homozygosity and providing proof-of-principle of the methods applied (see Appendix 5 in supplemental material). Inbred lines do not show indications of genetic load for traits related to reproduction and survival (see Appendix 2 in supporting information). These are the *non-evolving kin lines* abbreviated as ISO. By crossing these five different isofemale strains, we created an *evolving kin line* containing substantial genetic variation, further abbreviated as MIX. This was done by reciprocally crossing males and females of each of the isofemale strains: for each combination of strains, one female (last moulting stage) of strain X/Y was put together on a bean patch with three males of strain Y/X, allowing fertilisation (in case a fertilisation was unsuccessful, this step was repeated). The resulting F1, F2, and F3 generations were again mixed in such a manner that, eventually, we obtained one mixed strain (MIX) that comprised a mixture of all isofemale strains. Stock populations of the LS-VL and MIX strain were maintained on whole common bean plants in a climate-controlled room (28.1°C ± 2.1°C) with a light regime of 16:8 LD, while stock populations of the ISO strains were maintained on bean leaf rectangles in separate, isolated incubators (28°C, 16:8 LD). Before using the mite strains to initialise the experimental metapopulations, they were first synchronised. For each strain, sixty adult females were collected from the respective stock populations, placed individually on a leaf rectangle of 3.5 by 4.5 cm, and put in an incubator (30°C, 16:8 LD). The females were subsequently allowed to lay eggs for 24 hours, after which they were removed and their eggs were left to develop. Freshly mated females that had reached the adult stage one day prior to mating of the F1 generation were then used to initialise the experimental metapopulations (see below). As all mites were kept under common conditions during this one generation of synchronisation, direct environmental and maternally-induced environmental effects (Macke *et al.* 2011) of the stock conditions were standardised.

### Experimental range expansion

An experimental range expansion consisted of a linear system of populations: bean leaf squares (2 × 2 cm) connected by parafilm bridges [8 × 1 cm], placed on top of moist cotton. A metapopulation was initialised by placing ten freshly mated one-day-old adult females on the first patch (population) of this system. At this point, the metapopulation comprised only four patches. The initial population of ten females was subsequently left to settle, grow, and progressively colonise the next patch(es) in the linear array through ambulatory dispersal. Three times a week, all patches were checked and one/two new patches were added to the system if mites had reached the second-to-last/last patch. Mites were therefore not hindered in their dispersal attempts, allowing for a continuous expansion of the range. A regular food supply was secured for all populations by renewing all leaf squares in the metapopulation once every week; all one-week-old leaf squares were shifted aside, replacing the two-week-old squares that were put there the week before, and in their turn replaced by fresh patches. As the old patches slightly overlapped the new, mites could freely move to these new patches. Mites were left in this experimental metapopulation for approximately ten generations (80 days) during which they could gradually expand their range.

### Treatments

We performed two experiments, each of which contrasted two types of experimental metapopulations (see Figure 1). In the first experiment, we contrasted *unmanipulated* control LS-VL strains, further abbreviated as CONTROL, with a treatment where females in the metapopulations were replaced on a weekly basis by randomly chosen, but similarly aged, females from the LS-VL stock. This *Replacement From Stock* treatment is further abbreviated as REPLACEMENT. The metapopulations within the CONTROL treatment thus started with a high enough amount of standing genetic variation for evolution to act on. Kin structure was not manipulated in this treatment and kin competition was therefore expected to increase towards the range edge (see introduction). The REPLACEMENT treatment maintains age and population structure (i.e., if x females were on a patch before the replacement, they were replaced by x females from the stock) but prevented genetic sorting, and destroyed local relatedness, thus preventing both spatial sorting and kin competition. In this experiment, we thus compared unmanipulated, genetically diverse metapopulations (CONTROL treatment) with regularly reshuffled metapopulations in which only effects of density-dependent dispersal remained (REPLACEMENT treatment; cf. Ochocki *et al.* 2017). Both treatments were replicated six times.

**Fig 1.**
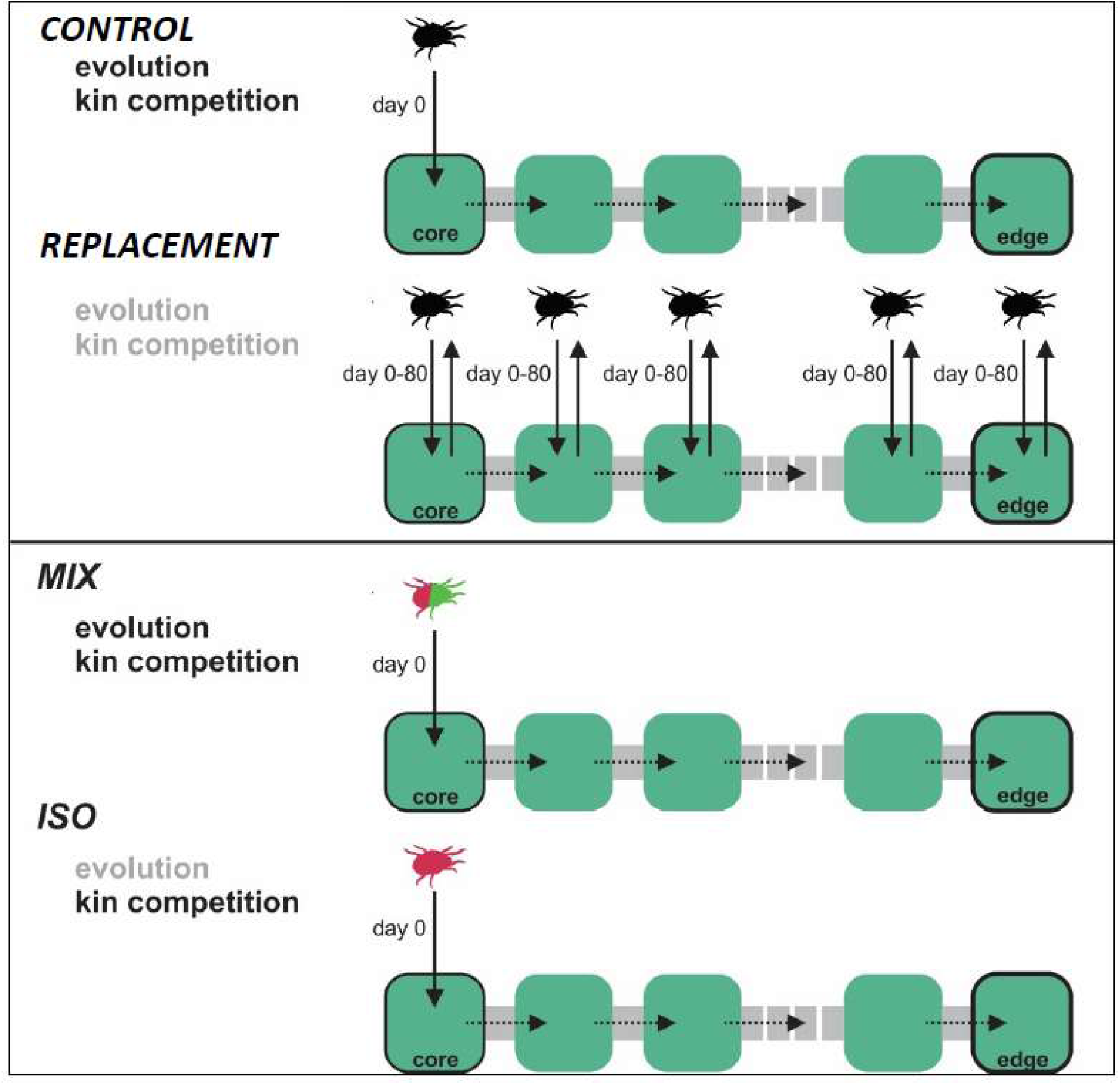
*The four treatments of the two experimental range expansion experiments. In the first experiment (upper panels), microcosms were either assigned to CONTROL (non-manipulated) or to REPLACEMENT (replacement from stock). Both these treatments harboured a relatively high amount of standing genetic variation (mites from LS-VL strain), but in REPLACEMENT all adult females were regularly replaced by females from the LS-VL stock while this was not the case for CONTROL. In the second experiment (lower panels), microcosms were either assigned to an evolving kin (MIX) or non-evolving kin line (ISO). The former harboured standing genetic variation (mites from MIX strain; different isofemale lines represented by a single mite colour), but the latter did not (here only one setup with a single isofemale line represented). No reshuffling was performed in this second experiment. By the end of the experiment, the final range size was measured as the number of occupied patches (dashed line thus denotes variable number of patches between the fixed core and emerging edge patch).*

In the second experiment, we compared *non-evolving kin lines* with *their evolving kin mixtures* (ISO versus MIX). Isofemale lines have been used in previous studies as an approach to have evolution-free controls in experimental evolution trials (Turcotte *et al.* 2011; Ochocki *et al.* 2017; Wagner *et al.* 2017). The evolving kin lines (MIX) harbour high levels of standing genetic variation. No manipulations of kin structure were performed. Experimental metapopulations with non-evolving kin lines (ISO) were initialised using mites from the SR-VP, JPS or LONDON isofemale strains. These metapopulations therefore harboured only a very limited level of standing genetic variation for evolution to act on (see Appendix 5 in supplemental material). As in the MIX treatment, kin structure was not manipulated. In this second experiment, we thus compared unmanipulated metapopulations (MIX treatment) with metapopulations where only condition dependency (density-dependent dispersal and kin competition) played a role (ISO treatment) (cf. Wagner *et al.* 2017). Both treatments were replicated six times (in case of ISO, two replicates, i.e., experimental range expansions, per isofemale strain were set up). Because our experimental metapopulations were initialised with an already mixed strain instead of with separate strains that would hybridise after initialization, a “catapult effect” with subsequent faster range expansions in the MIX treatment was prevented (Wagner *et al.* 2017).

The experimental range expansions started with a limited number of founders (10 females), thereby mimicking ongoing range expansion of *T. urticae* along a linear patchy landscape. Each replicated population invaded its respective landscape for ten generations (spanning 80 days). Genetic trait variation as determined in common garden experiments did not differ among any of the lines, likely due to the dominance of plasticity [see APPENDIX 2 in Supporting Information]. Starting from the same levels of trait variation, MIX and CONTROL thus represent treatments where range expansions are determined by evolution and putative kin interactions, ISO represents a treatment with high kin structure but restricted evolution, and REPLACEMENT a treatment constraining both kin competition and evolution. In addition to monitoring range expansions along the linear system, we quantified life history trait variation and genetic variation in gene expression between core and edge populations at an unprecedented level of detail (APPENDIX 3, 4 in Supporting Information). All data were analysed using general(ized) linear models with proper error structure; the individual traits longevity, fecundity, sex ratio and survival were integrated into a simulated growth rate measure by means of bootstrapping within replicates. A detailed overview of all used statistical models can be found in APPENDIX 3.

## Results

The eventually reached range size is a measure of range expansion speed. Despite the incorporation of uncertainties regarding condition-dependent dispersal thresholds, our model (APPENDIX 1 in Supporting Information) predicted range expansions to proceed at a 28.7% slower mean rate when signatures of both kin competition and spatial sorting were removed, while expansion rates were only 4% slower when only spatial sorting was prevented, but kin competition was present (Fig. 2).

**Fig 2.**
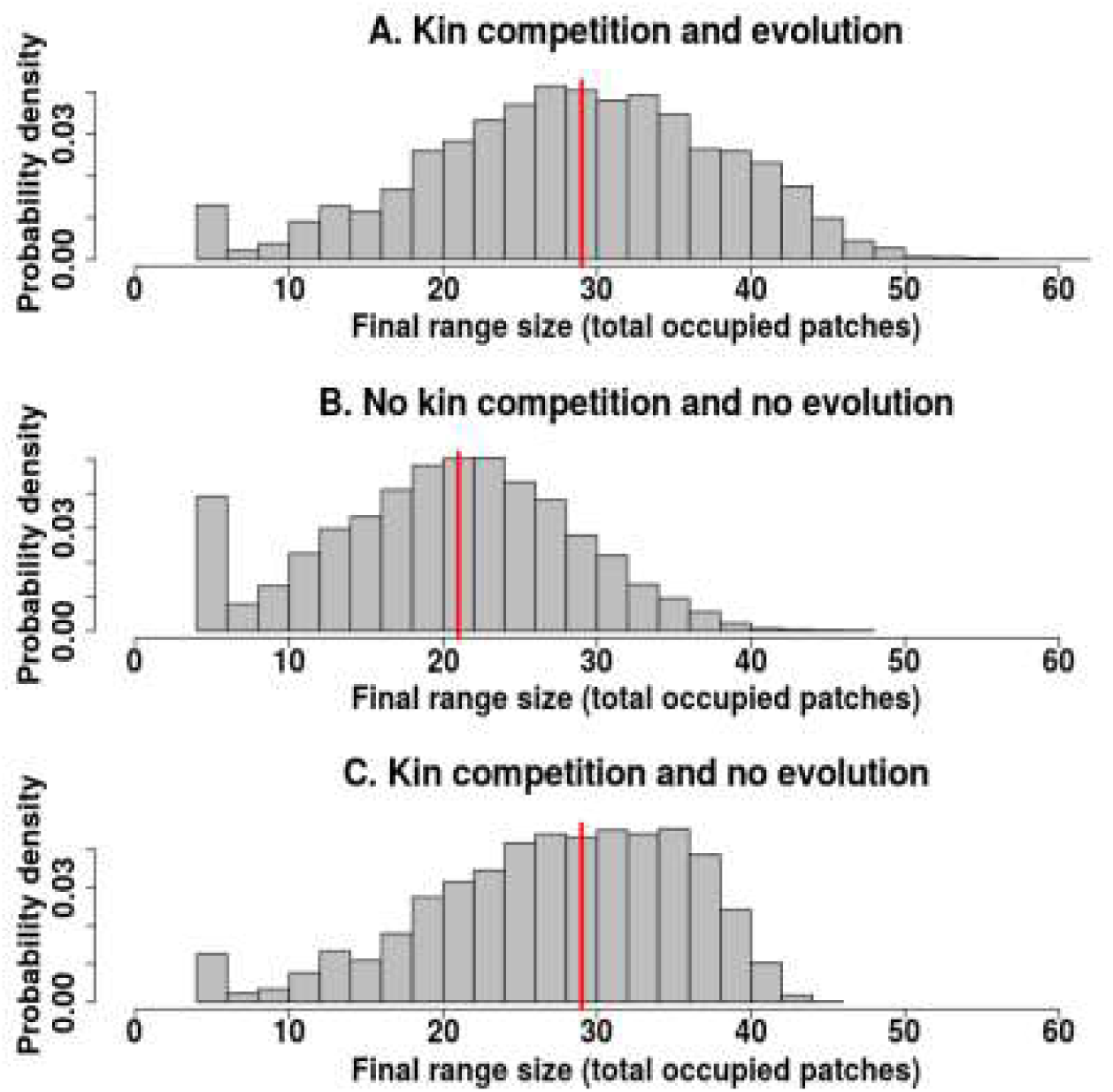
*Predicted range size (last patch occupied in the linear gradient) by a highly parameterised, stochastic model simulating expansion dynamics in the experimental setup over a period of 80 days (see appendix 1* in Supporting Information*). Overall, range expansions proceed more slowly when kin competition is excluded (B). Median values (indicated by the red lines) under the scenario with kin competition and spatial sorting (A) are similar to those for scenarios with kin competition but where spatial sorting is prevented (C). The scenario ‘No kin competition and spatial sorting’ was neither modelled in the individual-based models nor experimentally validated.*

In the experiments, we detected a 28% lower rate of range expansion in the treatment with mites replaced from stock, in which kin competition and evolution were constrained, versus the non-manipulated control treatment (CONTROL-REPLACEMENT contrast: GLM, day × treatment interaction for range size, F_1,54.8_=7.62; P=0.007; Fig. 3). However, no statistically significant differences were found in the experiment that contrasted the evolving with non-evolving kin lines that inhibited spatial sorting but left kin competition intact (MIX-ISO contrast: day × treatment interaction, F_1,71.1_=0.71; P=0.40). Differences in slopes were 0.082 ± 0.026 SE patches/day for the CONTROL-REPLACEMENT comparison and 0.030 ± 0.036 SE patches/day for the MIX-ISO comparison, with eventual range size matching the average model predictions. No significant differences in the variation in spread rate were present among any of the treatments (coefficients of variation in the reached distance with 95% CI based on bootstrapping: CONTROL: 0.246 [0.147-0.276]; REPLACEMENT: 0.2424 [0.133-0.279]; MIX: 0.279 [0.207-0.314]; INBRED: 0.220 [0.198-0.248]).

**Fig 3.**
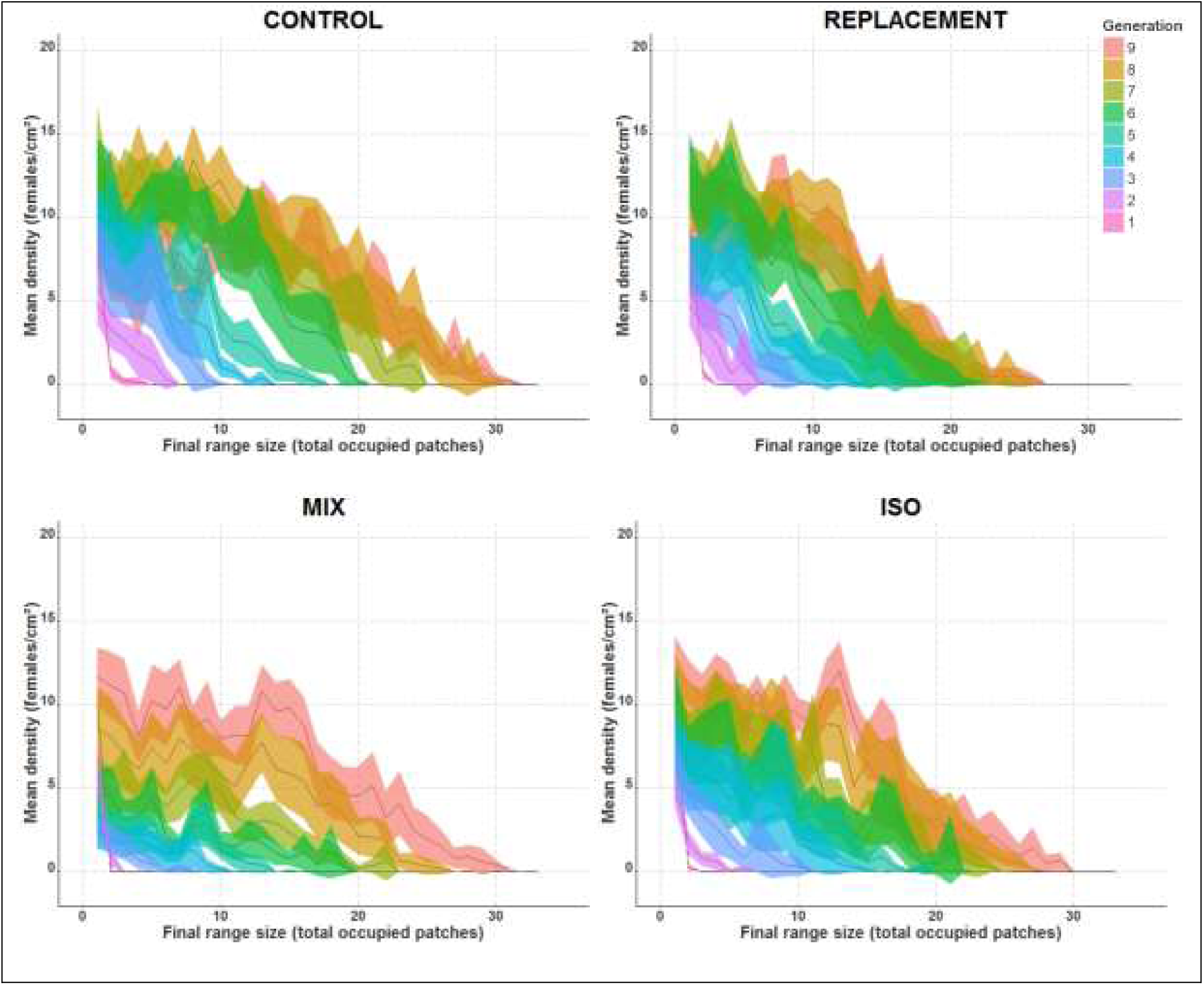
*Observed range expansion averaged (±STDEV: coloured belt) over the six replicates per treatment. Population densities are shown along the gradient from core to edge (distance). From generation to generation, the populations advanced their range (densities along the linear metapopulation).*

Furthermore, we tested whether increased range expansion resulted from evolved trait differences between edge and core populations [see APPENDIX 3 in Supporting Information]. In none of the treatments significantly higher dispersal rates were detected in individuals from the expanding front relative to the ones collected from the core patches. Therefore, the accelerated range advance in the treatments with unconstrained evolutionary dynamics was achieved independently of evolutionary changes in dispersiveness. Intrinsic growth rates, however, were systematically higher in edge relative to core populations in treatments that allowed evolutionary dynamics (CONTROL: F_1,153_=5.32, p=0.0225; MIX: F_1,235_=6.46, p=0.0117; See Fig. 4), but not in those where evolution was experimentally inhibited (REPLACEMENT: F_1,117_=0.31, p=0.582; ISO: F_1,216_=1.47, p=0.227; See Fig. 4). We found no indications of variation in any other fitness-and dispersal-related traits among the different treatments (see APPENDIX 3 in Supporting Information). We observed significant replica × location variation in traits during experimental evolution, eventually resulting in divergent trait covariances among replicates. Under such conditions, different life history strategies encompassing multivariate trait correlations but leading to similarly high population growth rates might eventually be spatially sorted.

**Fig 4.**
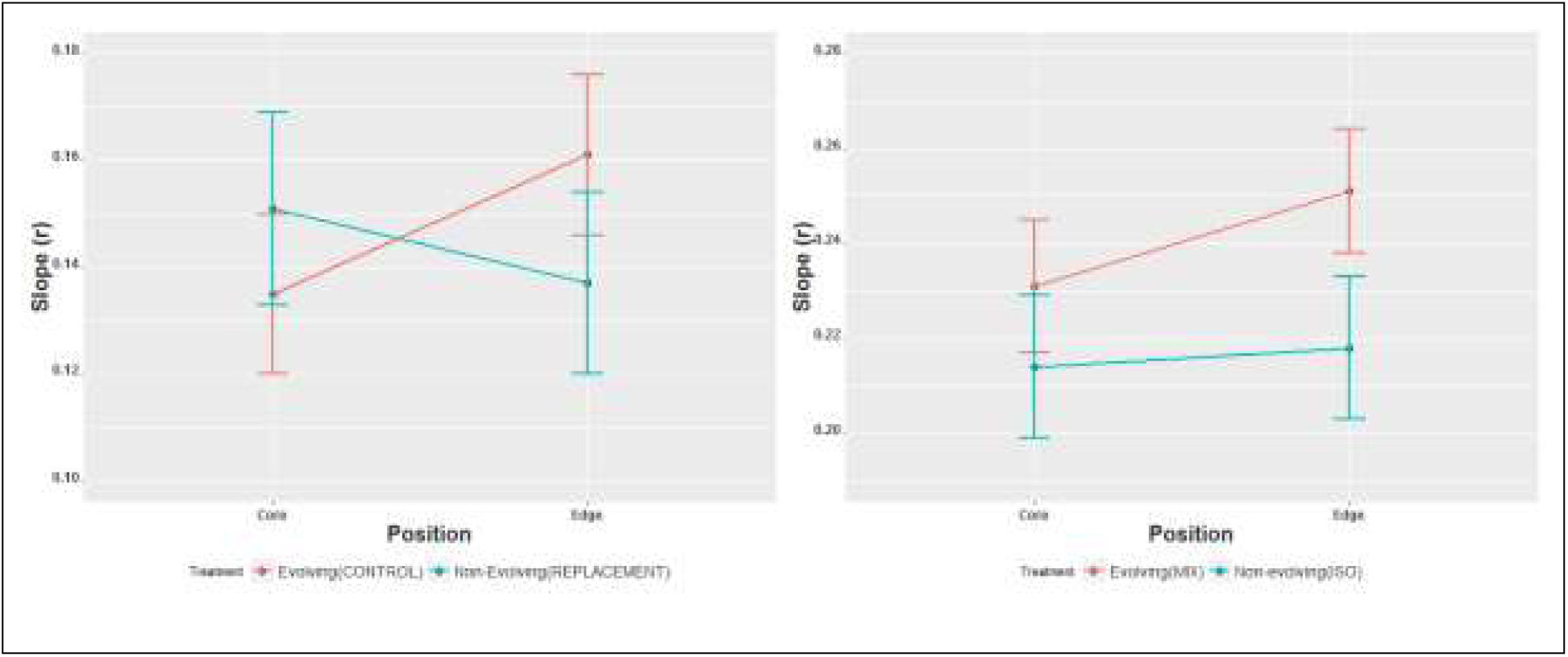
*Observed intrinsic growth rate (slope) expressed as the daily increase in Ln(population size) of offspring from a single female mite originating from the core versus edge in the two experimental range expansion contrasts. Treatments that allowed evolution (CONTROL, MIX) are red-coloured, those in which evolution was constrained (REPLACEMENT, ISO) are in blue*.

Bootstrapping growth rates did not follow the empirically determined higher growth rates at the leading edges (see APPENDIX 3 in Supporting Information). We attribute this opposing evidence from simulated relative to observed intrinsic growth rates either to the fact that the simulated ones systematically assume invariant life history trait expression during population growth, thereby neglecting density effects and other individual interactions or to the difficulty of detecting trait correlations at the individual level. We also neither found differences in quantitative genetic (APPENDIX 3 in Supporting Information) and transcriptomic (APPENDIX 4 in Supporting Information) trait variation between the inbred, mixed, and highly diverse stock populations, nor between core and edge populations, again indicating the dominance of plasticity over evolution for life-history traits in our model system.

## Discussion

Cooperation and conflict are central to understanding organismal interactions and their impact on population and community dynamics (West *et al.* 2002; Nadell *et al.* 2016). Dispersal, a central trait in life history (Bonte & Dahirel 2017) provides a prominent means to avoid competition with kin, thereby maximising long-term (inclusive) fitness. Dispersal also strongly impacts spatial dynamics in fragmented landscapes or during range expansions (Kubisch *et al.* 2014; Cheptou *et al.* 2017; Jacob *et al.* 2017). Despite the knowledge that genetic relatedness is increasing in edge populations during range expansions (Newman & Pilson 1997; Swaegers *et al.* 2013; Nadell *et al.* 2016), no connections have been made between relatedness-dependent dispersal as a behavioural response to kin competition and accelerating range expansions. So far, fast range expansions have been attributed to environmental changes (Keane & Smith 2002) and evolutionary processes (e.g., Phillips *et al.* 2010; Shine *et al.* 2011). We here present highly congruent results from experimental range expansions and a simple simulation model, showing that dispersal away from close relatives may be a key driver of fast range expansions. It thus provides an alternative but non-exclusive mechanism explaining fast range expansions despite potentially severe genetic diversity loss (Estoup *et al.* 2016).

Our experimental procedures enabled us to independently test the occurrence and importance of spatial sorting and relatedness (hence kin competition). This yielded several important insights. First, we demonstrate the presence of spatial sorting as an evolutionary process during range expansions. Evolutionarily increased growth rates at the leading expansion front are in good accordance with processes of spatial selection at the expanding front and are also in line with recent studies based on field observations (Phillips *et al.* 2010; Shine *et al.* 2011; Perkins *et al.* 2013), including earlier work on natural populations of the model species used here (Van Petegem *et al.* 2016a, b). Second, despite the evolutionary increase in growth rates at the expansion front, range expansions in the MIX treatment were on average not significantly faster than those from the genetically depleted lines (ISO). While both treatments maintained kin competition, kin competition was expected to be high throughout the range of the genetically depleted lines (ISO), while increasing in intensity to similar levels as in the ISO treatments at the advancing edge in the mixed lines. Spatial sorting in addition to normal levels of kin competition were thus approximately equivalent to high levels of kin competition without spatial evolution in determining the pace of our experimental range expansion. Third, and in line with our modeling results, we demonstrate by means of replacement procedures that disruptions of genetic structure–both in terms of genetic sorting and relatedness-substantially slow down range expansions. Given similar rates of evolutionary change in growth rates as in the MIX-ISO experiment, the CONTROL-REPLACEMENT experiment thus reinforces the idea that relatedness-dependent dispersal responses and spatial sorting are joint drivers of range expansions. Given the potential presence of drift and genetic load in the control treatments (although no evidence in this direction was found; see APPENDIX 3 in Supporting Information), our experimental range expansions even suggest that such behavioural responses may be more important than spatial selection, at least during the onset of the expansion. In line with the rather marginal importance of spatial sorting, we did not detect changes in spread rate variation. This contrasts with Williams *et al.* (2016a) where sorting narrowed variance in spread rate and the experiments by Ochocki *et al.* (2017) and Weiss-Lehman *et al.* (2017), which reported higher spread rate variance.

Our conclusion that increased relatedness at expanding range margins may further boost expansion dynamics is based on the assumption that organisms are able to assess local genetic relatedness via kin recognition abilities, and to exhibit an according conditional dispersal response. Kin recognition strategies can be categorised into (i) mechanisms based on predictable kin overlap in space and time (for example parent birds treating hatchlings in their nest as their offspring or larvae interacting with presumed relatives because eggs laid by parents are spatially concentrated), (ii) kin discrimination following initial interactions and learning (offspring getting habituated to cues from nestlings or individuals living nearby and learning to associate specific cues with presumed kinship), and (iii) kin discrimination based on innate, typically genetic cues that enable recognition of relatives (when kinds indicate kin; Queller 2011) under conditions that are not predictable in space or time (Waldman 1988). The first two mechanisms are obviously not relevant within our framework as they are independent of any spatial signature of genetic diversity loss and increased relatedness at expanding range margins. For relatedness-dependent dispersal to occur within a metapopulation or during range expansions, individuals need to be able to reliably assess the level of kin competition within a larger spatial neighbourhood, typically the local patch. Because gene flow renders levels of local relatedness dynamic, phenotype matching based on self-learning or recognition alleles is anticipated to be the main relevant mechanism of kin recognition and discrimination during range expansions. However, despite an increasing amount of empirical evidence in a wide array of species (e.g., Van Elsacker 1988; Tang-Martinez 2001; Biedrzycki *et al.* 2010; Le Vin *et al.* 2010; Bonadonna & Sanz-Aguilar 2012; Krause *et al.* 2012), support for the latter mechanism is much more controversial than for mechanisms related to learning or direct spatial associations (Gardner & West 2007). The current debate is strongly altruism-centred, fuelled by the absence of genetic kin-recognition strategies in many cooperating and/or group-living species. Within this context, theoretical developments demonstrate that genetic kin recognition mechanisms can only be stable when mutation rates are high, or when extrinsic mechanisms, such as parasite-host dynamics, maintain the diversity of alleles linked to cues involved in kin through fluctuating selection (Gardner & West 2007).

Dispersal decisions are taken hierarchically in response to multiple ultimate and proximate drivers (Matthysen 2012; Legrand *et al.* 2015). Because dispersal is theoretically leading to an ideal free distribution of fitness expectations in a metapopulation, it is a fitness-homogenising process (Bonte & Dahirel 2017). In response to local selection pressures, such as density, sex ratio, disturbance and relatedness (Bowler & Benton 2005), dispersal leads to the spread of genotypes in spatially structured systems. It is thus key to the maintenance of genetic diversity at the metapopulation level. Given that most likely all organisms inhabit spatially structured environments, kin(d)-recognition strategies might thus have primarily evolved to enable relatedness-dependent dispersal in order to avoid kin competition. This perspective would not only explain the existence of genetic kin recognition across a wide range of non-cooperative organisms, it also suggests that its link with range expansions is relevant far beyond our studied system. It is for instance not improbable that mobile vertebrates engage in exploratory behaviour to assess the level of relatedness with conspecifics in areas distant from the breeding territory (Delgado *et al.* 2014). Less mobile or sedentary sexual species (insects, plants) might by contrast use genetically based cues following in-and outbreeding to assess the level of relatedness at larger spatiotemporal scales from a set of close interactions with conspecifics.

If future research confirms the omnipresence of genetic kin(d)-recognition strategies (including in humans Wedekind *et al.* 1995; Jacob *et al.* 2002), spread mechanisms based on kin(d) recognition might be widespread, and, in organism with predominantly plastic reproduction and dispersal, potentially more relevant than the earlier proposed eco-evolutionary feedbacks following spatial sorting. From an even broader perspective, our work calls for a further integration of hitherto rather isolated disciplines related to the evolution of sociality and spatial dynamics to increase our understanding of patterns of biodiversity from local to regional scales.

## Acknowledgements

This project was funded by the Research Foundation – Flanders (FWO; project G.0610.11, G.018017N and G053815N). D.B. and R.S. were supported by the FWO research network EVENET; RS by the KU Leuven (PF/2010/07 and C16/17/002). Services used in this work were provided by the Flemish Supercomputer Center (VSC), funded by Ghent University, the Hercules Foundation, and the Flemish government (Department of Economy, Science, and Innovation). We are grateful to Jonathan Levine, Tom Miller and three anonymous reviewers for constructive comments that improved the paper. We thank Peter Demaeght, Sabina Bajda, Astrid Bryon and Wannes Dermauw for inbreeding spider mite lines and Simon Snoeck for genomic analysis.

